# Expression and purification of Protease Activated Receptor 4 (PAR4) and analysis with histidine hydrogen deuterium exchange

**DOI:** 10.1101/804427

**Authors:** Maria de la Fuente, Xu Han, Masaru Miyagi, Marvin T. Nieman

**Affiliations:** Department of Pharmacology, Case Western Reserve University, Cleveland, OH, USA

**Author notes:** **Correspondence:** Marvin T. Nieman, Department of Pharmacology, Case Western Reserve University, 2109 Adelbert Rd. W308B, Cleveland, OH, 44106-4965, USA.

## Abstract

Protease activated receptors (PARs) are G-protein coupled receptors (GPCRs) that are activated by proteolyis of the N-terminus, which exposes a tethered ligand that interacts with the receptor. Numerous studies have focused on the signaling pathways mediated by PARs. However, the structural basis for initiation of these pathways is unknown. Here, we describe a strategy for the expression and purification of PAR4. This is the first PAR family member to be isolated without stabilizing modifications for biophysical studies. We monitored PAR4 activation with histidine-hydrogen deuterium exchange (His-HDX). PAR4 has 9 histidines that are spaced throughout the protein allowing a global view of solvent accessible and non-accessible regions. Peptides containing each of the 9 His residues were used to determine the *t*_1/2_ for each His residue in apo or thrombin activated PAR4. The thrombin cleaved PAR4 had a 2-fold increase (p > 0.01) in *t*_1/2_ values observed for four histidine residues (His_180_, His_229_, His_240_, and His_380_) demonstrating that these regions have decreased solvent accessibility upon thrombin treatment. In agreement, thrombin cleaved PAR4 also was resistant to thermolysin digestion. In contrast, activation with the PAR4 agonist peptide was digested at the same rate as apo PAR4. Further analysis showed the C-terminus is protected in thrombin activated PAR4 compared to uncleaved or agonist peptide treated PAR4. The studies described here are the first to examine the tethered ligand activation mechanism for a PAR family member using biophysically and shed light on the overall conformational changes that follow activation of PARs by a protease.

## INTRODUCTION

Protease activated receptors (PARs) are the primary means by which proteases directly initiate intracellular signaling.^1^ There are 4 PARs (PAR1-4) that comprise a specialized family of G-protein coupled receptors (GPCRs), which have a unique activation mechanism whereby cleavage of the N-terminus generates a tethered ligand. PAR1 was originally identified as the thrombin receptor on platelets, however it is now recognized that PARs are expressed in many cell types and are activated by a growing panel of proteases.^2,3^ Once activated, PARs signal through Gα_q_, Gα_12/13_, Gα_i_, and arrestin depending on the activating protease and cellular context.^3^ It is reasonable to infer that these divergent signaling events are due to allosteric modulation of PARs by cofactors or specific tethered ligands. However, this has not been tested formally.

PAR4 was identified seven years after PAR1 and, due to its kinetics of activation, was initially considered a low affinity backup thrombin receptor on human platelets.^4^ The initial characterization of PAR4 revealed that the N-terminus of PAR4 does not have a hirudin-like sequence, which dramatically lowers the efficiency of PAR4 cleavage by thrombin.^5–9^ The net result is a ∼10-fold higher threshold for thrombin activation of PAR4 compared to PAR1. Once activated, PAR4 mediates prolonged signaling in platelets, which is required for sustained platelet activation and stable thrombus formation.^10^ In contrast PAR1 elicits a rapid, short-lived signal.^11,12^ Combining these observations, PAR4 has emerged as a therapeutic target for antiplatelet agents that prevent sustained activation by high levels of thrombin that are generated at the site of injury, while sparing PAR1 signaling.^13,14^ Initial clinical studies suggest that this therapeutic approach allows the formation of the initial hemostatic plug while preventing unwanted thrombosis.^14,15^

The activation of PARs via the tethered ligand mechanism was described in 1991 for PAR1.^2^ To date, there is still limited structural information regarding this unique mode of activation. High resolution crystal structures of PAR1 and PAR2 bound to antagonists have been solved.^16,17^ These structural studies reveal the molecular details of the receptor/antagonist interaction, but provide limited information on the tethered ligand mechanism due to modifications required to facilitate crystallization.^18^ Specifically, the N-and C-termini were removed to limit overall flexibility, and T4 lysozyme (PAR1) or BRIL (PAR2) was inserted into intracellular loop 3 (ICL3) to further stabilize the protein.^16^ For PAR2, additional stabilizing mutations were introduced.^17^ To date, there is no structural information for PAR4. A major roadblock in understanding the structural basis for the tethered ligand-mediated activation of PARs is the lack of structural studies with the native protein to allow the identification of the ligand binding site and the structural dynamics specific to this activation mechanism. Advances in mass spectrometry now allow protein dynamics to be monitored in response to activation or ligand binding. Histidine Hydrogen-Deuterium Exchange (His-HDX) takes advantage of the slow exchange rate of the imidazole C2-hydrogen on the histidine residue.^19–21^ This is in contrast to amide-HDX, which measures the exchange rate of backbone hydrogens. The histidine residues are effectively site-specific probes that monitor changes in the molecular environment at these sites as the protein changes conformation upon activation or ligand binding. PAR4 has nine histidine residues that are spaced throughout the protein, which allows an overall evaluation of the conformational changes using His-HDX. In addition, the slow back exchange allows analysis of proteins in complex mixtures.^19,21^

Here, we used histidine hydrogen deuterium exchange (His-HDX) to examine the conformational changes that occur within PAR4 upon activation by thrombin or the PAR4 activation peptide AYPGKF. Taken together, our results demonstrate that cleavage of PAR4 by thrombin leads to a dramatic rearrangement, which results in a more stable structure with increased resistance to experimental proteolysis. In contrast, the structural stability of PAR4 was unchanged following stimulation by AYPGKF compared to the non-activated receptor. These studies are first to examine the structural rearrangement of PAR4 upon activation and may provide a molecular basis for the differential signaling that is observed from PAR activation peptides.^22^

## MATERIALS AND METHODS

### Reagents and Antibodies

Monoclonal antibodies to PAR4, 14H6 and 2D6, were produced in our laboratory and have been described.^23^ The goat polyclonal antibody PAR4-C20 (catalog # SC-8464, lot # B1513) was purchased from Santa Cruz Biotechnology (Dallas, Texas). Rabbit polyclonal antibodies to G_q/11_α (catalog # 06-709, lot # 2952616) and Gβ (catalog # 06-238, lot # 2836919) were acquired from EMD Millipore (Temecula, California). The monoclonal antibody to polyHistidine, Clone HIS-1 (catalog # H1029, lot # 106M4768V), was purchased from Sigma-Aldrich (St. Louis, MO). The hybridoma for the monoclonal antibody, FLAG M1 (clone 4E11), was purchased from American Type Culture Collection (ATCC). The rhodopsin monoclonal antibody, 1D4, was kindly provided by Dr. Vera Moiseenkova-Bell.

Human α-thrombin (catalog # HCT-0020, specific activity greater than 2989 U/mg) was purchased from Hematological Technologies (Essex Junction, VT). Trypsin (catalog # V511A, lot # 0000079904) and thermolysin (catalog # V400A, lot # 0000142688) were bought from Promega (Fitchburg, WI). TLCK treated chymotrypsin (catalog # LS001432) was from Worthington (Lakewood, NJ). FLAG (DYKDDDDA) and 1D4 (TETSQVAPA) were synthesized at GenScript (Piscataway, NJ). The PAR4 activation peptide, AYPGKF, was purchased from Tocris Bioscience (Minneapolis, MN). N-Dodecyl-β-D-Maltopyranoside (DDM) and Lauryl Maltose Neopentyl Glycol (NG310, LMNG) were from Anatrace (Maumee, OH). All other reagents were from Thermo Fisher Scientific (Pittsburgh, PA) except where noted.

The 10% acrylamide gels used for in gel-extraction and digestion were pre-casted gels from Bio-Rad (Hercules, CA). Native gels, buffers, reagents, and ladder were purchased from Novex by Life Technologies-Thermo Fisher Scientific (Waltham, MA), and were used following the manufacturer’s protocols. All other gels used for Coomassie stain or Western Blots were casted in the lab using ProtoGel 30% from National Diagnostics (Atlanta, GA). Deuterium oxide (D2O, 99%) was purchased from Cambridge Isotope laboratories (Andover, MA).

### Cloning and Cell Culture

The cDNA for BRIL-PAR4 was codon optimized for expression in Sf9 cells and synthesized by GenScript. The cDNA for Gαq (catalog # GNA0Q00000) was purchased from cDNA Resource Center (Bloomsburg, PA). The cDNA for Gβ1 (catalog # 67016) and Gγ2 (catalog # 67018) were purchased from Addgene (Cambridge, MA). All constructs were cloned into pFastBac and expressed in Sf9 cells using the Baculovirus Expression Vector System (Invitrogen, Carlsbad, CA) grown in ESF 921 media from Expression Systems (Davis, California). The recombinant bacmids were transfected into Sf9 cells using Cellfectin II Reagent for production of baculovirus for BRIL-PAR4, Gαq, Gβ1, or Gγ2 that was subsequently used for protein expression in Sf9 cells using manufacturer’s protocol (Invitrogen).

To map the C-terminal epitope of the C-20 antibody, we transiently expressed wild type PAR4 and two C-terminal truncation mutants at amino acid 358 (RAGLFQR*) or 373 (KASAEGG*) in HEK293 cells.^24^ Transfections were done using Lipofectamine 2000 from Thermo Fisher Scientific (catalog # 11668019) following manufacturer’s protocol. Cells were harvested at 48 hr post transfection and lysed in RIPA buffer.

### Protein Purification

Membranes of Sf9 cells expressing BRIL-PAR4 were prepared by resuspending cell pellets in lysis buffer (25 mM Tris pH 8, 300 mM sucrose, and 5 mM EDTA plus protease inhibitors (cOmplete tablets-EDTA free by Roche-Basel, Switzerland). Cell suspensions were lysed in a microfluidizer followed by centrifugation at 3000*g* for 10 min and 14,000*g* for 45 min to remove cell debris. Finally, membranes were pelleted at 100,000*g* for 60 min. All centrifugations were done at 4 °C. Once the membranes were obtained, they were solubilized at 4 °C for 2 hours in 0.5% DDM, 20 mM HEPES pH 7.5, 50 mM NaCl, 0.1% Cholesteryl hemisuccinate (Sigma, St. Louis, MO), 20% glycerol and 1 mM DTT, plus proteinase inhibitors. The BRIL-PAR4 for Histidine Hydrogen-Deuterium exchange was purified by taking advantage of the C-terminal 1D4 epitope (Figure 1) for affinity chromatography with 1D4-coupled sepharose beads. Prior to elution, the buffer was exchanged to 0.1% LMNG, 20mM HEPES pH7.5, 50mM NaCl, 0.02% Cholesteryl hemisuccinate, 5% glycerol and 1 mM DTT. The column was eluted using a peptide corresponding to the 1D4 epitope. For thermolysin digestions, PAR4 was purified by using FLAG-M1 coupled to CNBr-activated sepharose (GE Healthcare, Chicago IL) adding 2 mM CaCl_2_, and 150 mM NaCl, to the column buffer of 0.1% DDM, 20mM HEPES pH7.5, CHS 0.02%, 5% Glycerol, and 1 mM DTT.. The columns were eluted with FLAG peptide (DYKDDDDA) and 5 mM EDTA. The affinity purified BRIL-PAR4 was digested with TEV to separate BRIL from PAR4 (see Figure 1). The PAR4 was concentrated using Vivaspin 2 Concentrators 50,000 MWCO PES, filtered with ultra-free centrifugal filter units 0.65um DV durapore (Millipore), and injected in a FPLC column Superdex 200 Increase 10/300GL. The final concentration was ∼1 mg/ml.

**Figure 1.**
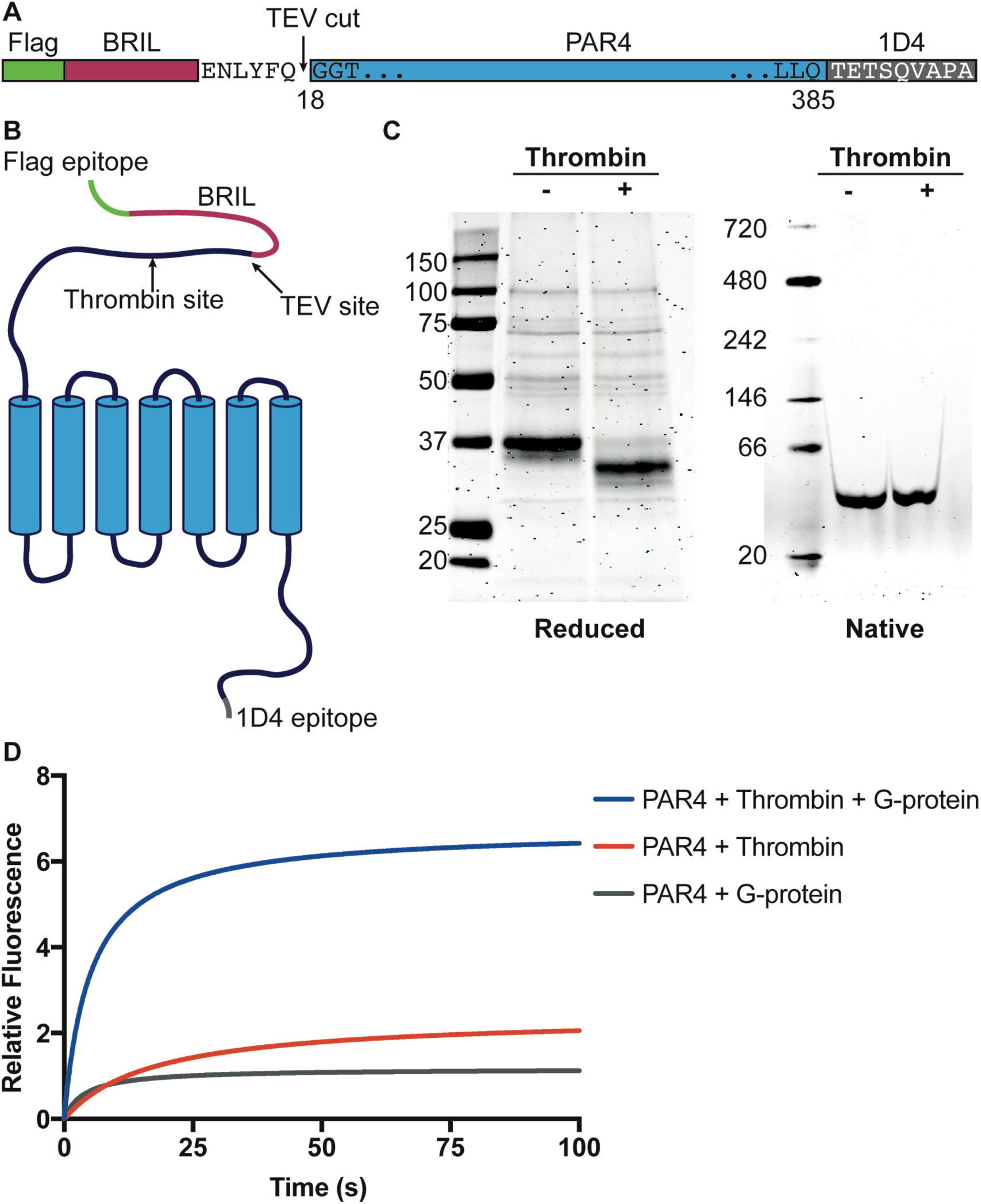
Expression and purification of human PAR4 from Sf9 insect cells. Linear diagram illustrates human PAR4 construct for expression in *Sf9* cells. Full-length human PAR4 was expressed as a BRIL-fusion protein with an N-terminus Flag tag and a C-terminus 1D4 tag (A). Representation of PAR4 (blue) fused with BRIL (red) with two epitope tags on both N- and C-termini. Thrombin and TEV cleavage sites are both marked on the diagram (B). The purity and quantity of purified PAR4 was confirmed with Coomassie blue stained SDS-PAGE (left) and native-PAGE (right). Thrombin cleavage results a 2.9 kDa shift of PAR4 from ∼37 kDa to ∼34 kDa on SDS-PAGE (C). G-protein activation assay that monitors relative fluorescence change of an intrinsic tryptophan confirms that the purified PAR4 is fully functional (D).

The Gαq and Gβ1 proteins were expressed without tags; the Gγ2 was expressed with a 6-His tag for purification. Baculovirus preps for each subunit (Gαq, Gβ1, and Gγ2) were combined in an optimized ratio to produce the Gαβγ complex in Sf9 cells. Cell membranes were prepared as described above for BRIL-PAR4. Solubilization of membrane preps was done at 4 °C overnight in buffer containing: 0.1% DDM, 1.5% Na Cholate, 20mM HEPES, 100mM NaCl, 5mM MgCl_2_, 50uM GDP (catalogue #G7127) Sigma Aldrich, 1mM DTT, and protease inhibitors, pH 7.5. The Gαβγ complex was purified with a Ni-NTA agarose column (Catalogue # 30210) from Qiagen (Hilden, Germany) followed by size exclusion chromatography in 0.02% DDM, 20mM HEPES, 100mM NaCl, 5mM MgCl_2_, 40uM GDP, 5 % Glycerol and 1mM DTT, pH 7.5 to separate dissociated Gβγ from the trimer.

### Histidine Hydrogen Deuterium Exchange (His-HDX) coupled with Mass Sepctrometry

Purified recombinant PAR4 was analyzed with His-HDX as previously described.^19,21^ Briefly, apo or thrombin cleaved PAR4 were incubated in 0.1% LMNG, 20mM HEPES pH7.5 made with D_2_O containing 50mM NaCl, 0.02% Cholesteryl hemisuccinate, 5% glycerol and 1 mM DTT at 37 °C for 72 hr. The samples were run on SDS-PAGE followed by staining with Coomassie blue. The electrophoresis and the subsequent staining were carried out at 4 °C to minimize the deuterium to protium back exchange. The bands corresponding to PAR4 were excised, cut into small pieces, and divided into 2 tubes. The samples were processed for in gel digestion with trypsin or chymotrypsin for 1 hr at RT. Following digestion, the trypsin and chymotrypsin digests were pooled, dried in a Speed Vac concentrator, redissolved in 0.1% formic acid, and then analyzed by LC/MS/MS using LTQ-Orbitrap XL mass spectrometer as described previously.^20^ The peptides were resolved and the half-life (*t*_½_) for H/D exchange was calculated as previously described.^19–21^ The data are from three independent experiments.

### Limited Proteolysis with thermolysin

Purified recombinant apo-PAR4 or thrombin cleaved PAR4 samples were treated with thermolysin (0.01 µg/reaction) for 0, 2, 5, 10, 20, 60 or 90 minutes, reactions were stopped with the addition of 50 mM EDTA and Laemmeli sample buffer containing 2-mercaptoethanol (5%). Samples were run on 12% SDS-PAGE and stained with QC Colloidal Coomassie blue (BioRad, catalog # 161-0803)). The rate of thermolysin digestion of PAR4 was quantified using the Li-Cor Odyssey imaging system (Li-Cor, Lincoln, NE). To follow the stability of the N- and C-termini, thermolysin digestion was followed by Western blotting using the antibody 2D6 to monitor the N-terminus and the antibody C-20 to monitor the C-terminus.

### G protein activation assay

The functionality of PAR4 was verified with a G protein activation assay that monitors the increase of intrinsic tryptophan fluorescence of the Gα subunit as the trimeric complex dissociates to Gαq and Gβγ.25 The fluorescence increase was monitored at 300 nm excitation and 335 nm emission at 20 °C. The reaction contained PAR4 (50 nM), Gαqβγ (500 nM), and GTPγS (Catalogue # 10220647001) Roche, (Basal, Switzerland) (300 µM) and was initiated with the addition of thrombin (100 nM). Fluorescent data was acquired for 500 sec. Control experiments were done as above without Gαqβγ complex or without thrombin.

## RESULTS

### Expression and purification of human PAR4

The goal of this study was to examine the structural rearrangements of PAR4 following activation by thrombin using histidine hydrogen deuterium exchange (His-HDX). Since we were interested in the conformational changes of PAR4 as a result of activation, the recombinant PAR4 was free of stabilizing modifications that are typical for structural studies with GPCRs (Figure 1).^18,26,27^ Human PAR4 was expressed as a apocytochrome *b*562 (BRIL) fusion protein with epitope tags on the N- and C-termini (Figure 1A and B). The addition of BRIL to the N-terminus of PAR4 increased expression >20-fold over our original constructs that did not contain it (data not shown). The signal sequence from the influenza hemagglutinin is removed during trafficking and processing the recombinant PAR4 in the *Sf9* cells leaving an N-terminal FLAG tag that was used for affinity purification.^28^ The 1D4 antibody epitope (TETSQVAPA) was added to the C-terminus to facilitate a second affinity purification strategy. Following purification, FLAG and BRIL were removed from PAR4 with tobacco etch virus (TEV) protease. The TEV recognition sequence was engineered such that the Gly residue at the P1 site was incorporated into the PAR4 sequence as Gly^18^; the first amino acid of the mature PAR4 protein (Figure 1A).^4^ The final purified recombinant human PAR4 migrated as a single band at the expected 37 kDa on reduced SDS-PAGE as indicated by Coomassie Blue staining (Figure 1C). Thrombin removed 2.9 kDa (Gly_18_-Arg_47_) and the resulting activated PAR4 migrated at the expected 34 kDa (Figure 1C). Native SDS-PAGE confirmed that the purified PAR4 did not form aggregates following purification (Figure 1C). The functionality of the purified PAR4 was confirmed with an *in vitro* G-protein activation assay that monitors the increase in intrinsic protein fluorescence of the Gα subunit.^25^ Stimulation of recombinant PAR4 with thrombin (100 nM) resulted in a rapid increase in fluorescence indicating activation of the G-protein complex (Figure 1D). Importantly, there was no increase in fluorescence in the absence of the Gαβγ complex or absence of thrombin (Figure 1D).

### Histidine Hydrogen-Deuterium exchange (His-HDX)

His-HDX monitors the rates of hydrogen-deuterium exchange at the C_2_ position on the imidazole ring of histidine residues (Supplemental Figure 1A).^19–21^ The exchange is slow at this position, allowing for long time scale measurements using histidine residues as site-specific probes. PAR4 has 9 histidine residues that are well spaced throughout the protein (Figure 2A), which provides a global view of solvent accessible and non-accessible regions. Following incubation in D_2_O for 72 h, unique peptides that contained each of the 9 histidine residues were generated by trypsin and chymotrypsin digestion (Supplemental Table 1). These peptides were used to determine the exchange rate (*t*_½_) for each histidine residue in non-activated or thrombin activated PAR4 (Table 1 and Figure 2B). The histidine exchange rates for PAR4 can be categorized into three groups; 1) a long half-life that did not change upon activation, 2) a moderate half-life that did not change upon activation, and 3) a moderate half-life that was prolonged upon PAR4 activation. The range of responses allows both a global view of PAR4 activation as well as site specific conformational changes.

**Table 1.** Exchange rates of histidine residues in PAR4.

| Table 1. Exchange rates of histidine residues in PAR4 |  |  |  |
| --- | --- | --- | --- |
| | $t_{1/2}$ (days) | | *p* value |
|  | Untreated | Thrombin |  |
| His<sub>136</sub> | 7.52 ± 1.07 | 8.88 ± 3.19 | 0.52 |
| His<sub>159</sub> | >40 | > 40 |  |
| His<sub>180</sub> | 8.42 ± 0.61 | 12.88 ± 2.37 | 0.01 |
| His<sub>229</sub> | 4.87 ± 0.02 | 9.25 ± 1.64 | 0.01 |
| His<sub>240</sub> | 4.13 ± 0.29 | 6.28 ± 0.43 | 0.01 |
| His<sub>269</sub> | > 40 | > 40 |  |
| His<sub>280</sub> | 10.61 ± 2.02 | 9.35 ± 0.54 | 0.44 |
| His<sub>306</sub> | 8.05 ± 2.00 | 10.59 ± 3.41 | 0.32 |
| His<sub>380</sub> | 3.11 ± 0.29 | 5.54 ± 0.55 | 0.003 |

**Figure 2.**
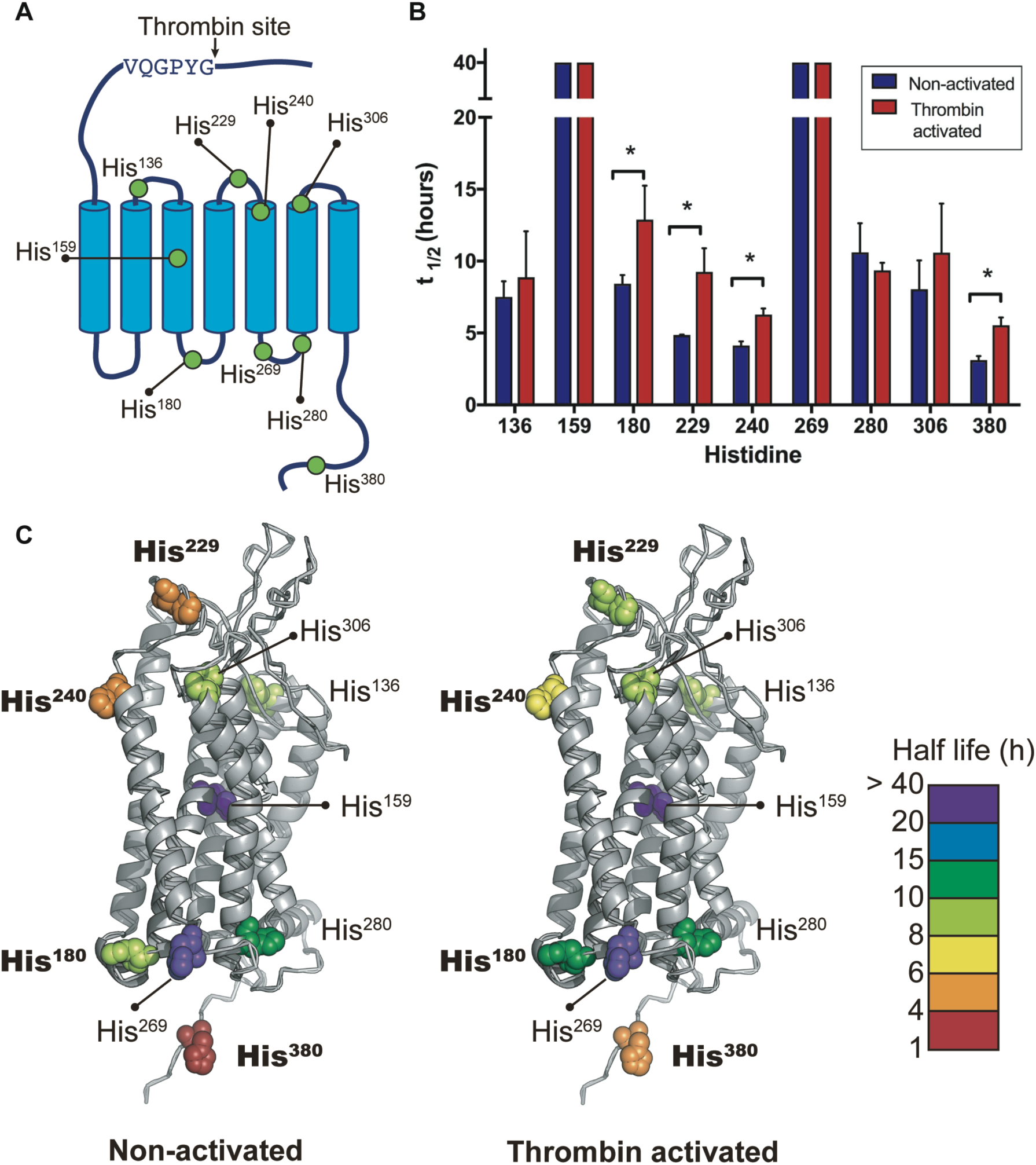
Distribution of histidine residues in PAR4 and representation of the exchange rates of histidines. The location of the 9 histidine residues are indicated on the diagram. These are distributed throughout the receptor and serve as probes to detect water accessibility in these regions (A). Bar graph showing the half-life (t_1/2_) for each histidine residue in non-activated or thrombin activated PAR4 (B) (n=3). PAR4 homology computational model for the comparison between the non-activated (apo state) versus the thrombin activated PAR4. Half-life of each histidine is represented by color coded scale (C).

The ligand binding site of PAR4 has not been determined experimentally. However, based on comparisons with PAR1, ECL2 is predicted to play a role.^16,29^ Four histidines (His_136_, His_229_, His_240_, and His_306_) are located in extracellular loops (ECL) of PAR4. His_136_ is located in ECL1 (shown in light green, Figure 2C) had a half-life of 7.52 ± 1.07 d in the non-activated state and did not change upon activation by thrombin (8.88 ± 3.19 d, *p* = 0.52) (Table 1 and Figure 2B). Similarly, His_306_, which is located in ECL3 (light green Figure 2C), also did not show a change in *t*_½_ upon activation, 8.05 ± 2.00 d versus 10.59 ± 3.41 d, *p* = 0.32. In contrast, His_229_ and His_240_, both located in ECL2, had a longer *t*_½_ upon activation. His_229_ has ∼2-fold shift from 4.87 ± 0.02 d to 9.25 ± 1.64 d (orange to light green, Figure 2C). His_240_ is also significantly prolonged from 4.13 ± 0.29 to 6.28 ± 0.43 (orange to yellow, Figure 2C). These data suggest that the N-terminal tethered ligand of PAR4 is interacting with ECL2 upon activation to mediate signaling.

The *t*_½_ values for His_159_ and His_269_ (shown in purple, Figure 2C) were >40 d in both states (non-activated and thrombin activated) indicating that these were not exposed to solvent and did not change in response to thrombin. The long half-life of His_159_ is expected as it is buried within transmembrane helix 3 (TM3) (Figure 2A and C). His_269_ is located at the base of TM5. Homology models with other GPCRs indicate that His_269_ is surrounded by other TM helices (Figure 2A and C), which likely contributes to its prolonged half-life in both states.

The cytoplasmic surface of GPCRs mediate signaling through interactions with G-proteins and arrestins. Two histidines, His_180_ and His_280_, are located in the intracellular loops (ICL) and His_380_ is located at the C-terminal tail. The *t*_½_ values for His_180_ increased from 8.42 ± 0.61 to 12.88 ± 2.37 d, *p* = 0.01 (Figure 2B) and indicated from light green to green in Figure 2C. In contrast, His_280_ did not change upon activation (Figure 2C, shown in green). His_380_ is located 6 amino acids from the C-terminus. The *t*_½_ nearly doubled from 3.11± 0.29 to 5.54 ± 0.55 d, *p* = 0.003 upon activation suggesting that this region of PAR4 dramatically changes following activation (Figure 2B and C).

In summary, the His-HDX experiments identified regions important for both receptor activation on the extracellular loops and signal transduction in the intracellular face. The increase in half-life for His_229_ and His_240_ demonstrates that ECL2 changes its molecular environment upon activation. A role for ECL2 in PAR4 activation is consistent with other PARs based on sequence homology. Finally, the His residues that have a longer half-life (*t*_½_) of H/D exchange following cleavage by thrombin are located throughout PAR4 indicating that PAR4 undergoes a global rearrangement and becomes more compact and potentially more rigid upon activation.

### Thrombin activated PAR4 is resistant to proteolysis

Resistance to proteolysis has long been used as a measure of structural stability.^30,31^ To validate the results from our studies using His-HDX, we compared the overall structural changes in PAR4 following activation with limited proteolysis by the low specificity metalloproteinase thermolysin. Initial experiments determined that 20 nM thermolysin was required to digest PAR4 to completion in 60 min (data not shown). We next analyzed the stability of PAR4 or thrombin activated PAR4 by monitoring the thermolysin reaction over time (1-90 min) via SDS-PAGE followed by Coomassie blue staining. A representative gel from three independent experiments is shown in Figure 3A. The gels where quantitated by measuring the disappearance of the two bands at ∼37 kDa using a Li-Cor Odyssey imaging system. Non-activated (apo) PAR4 was digested to completion in 60 min with a half-life (*t*_½_) of 11.9 ± 1.0 min and first order rate constant of 0.060 ± 0.005 min_-1_. The error is reported as the standard deviation from three independent experiments. Thrombin activated PAR4 did not show appreciable proteolysis at 90 min. Taken together, these data support the results from the His-HDX studies that PAR4 globally becomes structurally more compact upon cleavage by thrombin.

**Figure 3.**
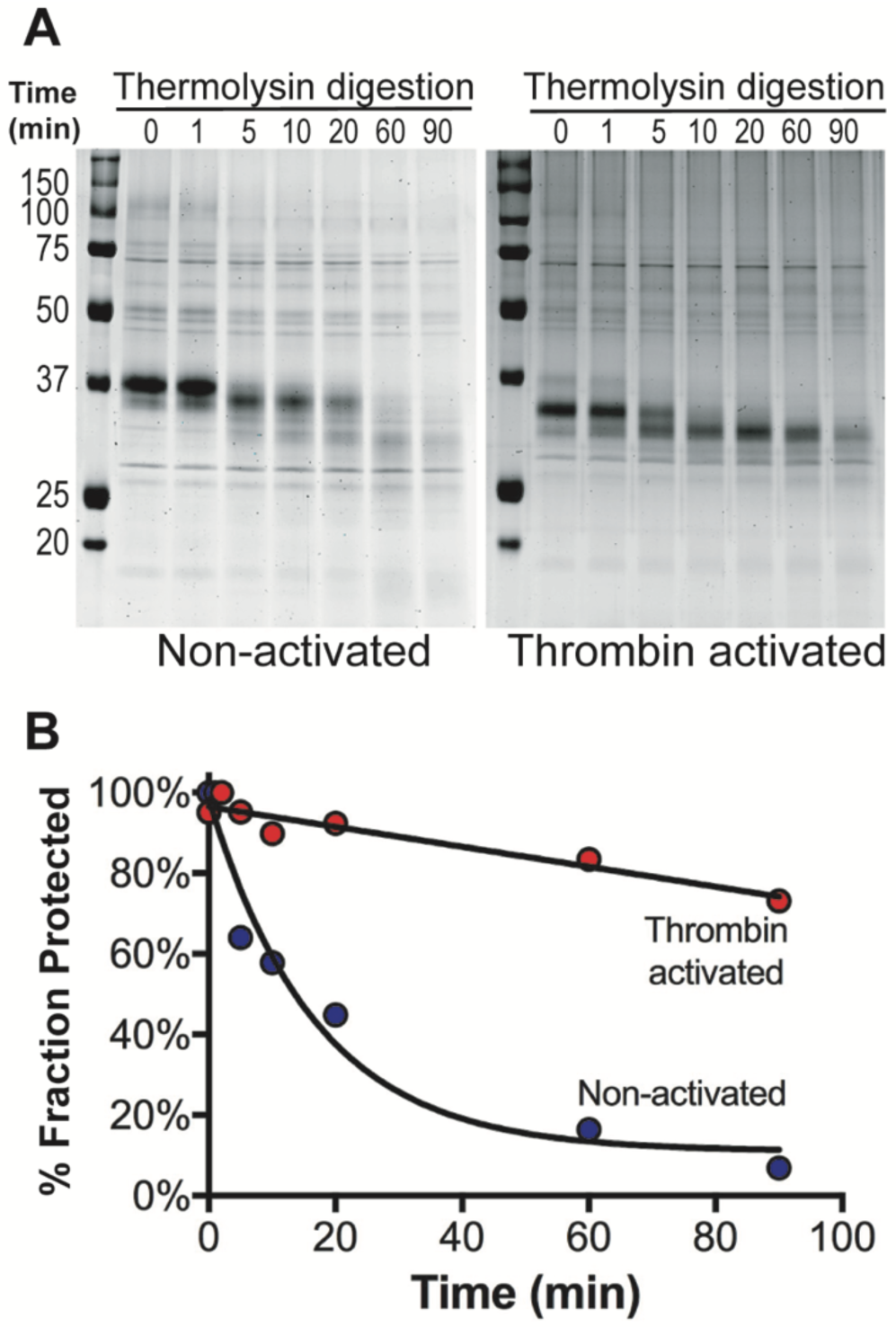
Stability of PAR4 monitored with proteolysis. Human PAR4 samples, full length and thrombin cleaved, were partially digested by thermolysin. Reactions were stopped at 1, 5, 10, 20, 60, or 90 min. Representative SDS-PAGE gels are shown in (A) and quantified in (B). (n=3)

### Mapping the proteolysis

The His-HDX and initial limited proteolysis experiments show that PAR4 becomes more compact and protected from proteolysis upon activation. We next determined which region of PAR4 was becoming protected by monitoring the thermolysin digest by Western blotting with antibodies directed to the N - or C-terminus of PAR4. The mouse monoclonal antibody 2D6 binds to the N-terminus at (G_74_WVPTR_78_), which is near the start of transmembrane helix 1 (**Supplemental Figure 3**).^23^ The goat polyclonal antibody (C20) maps to the last 12 amino acids of PAR4 (**Supplemental Figure 3**). These epitopes were monitored simultaneously using secondary antibodies labeled with distinct infrared probes and the Li-Cor Odyssey System. The 2D6 epitope was lost at 90 min in apo-PAR4 (Figure 4A, green). The C-terminus was also progressively cleaved to generate a profile of proteolytic products that reached a maximum at 120 min (Figure 4A, red). The 2D6 epitope was lost at 60 min for thrombin cleaved PAR4 indicating that the tethered ligand may lead to an exposed region in the N-terminus that is accessible to thermolysin. In contrast, the C-terminus was resistant to proteolysis indicating that it is less accessible to thermolysin. These data are in agreement with the His-HDX data showing that His_380_ has a longer t_1/2_ showing that it less accessible H/D exchange and protected from proteolysis.

**Figure 4.**
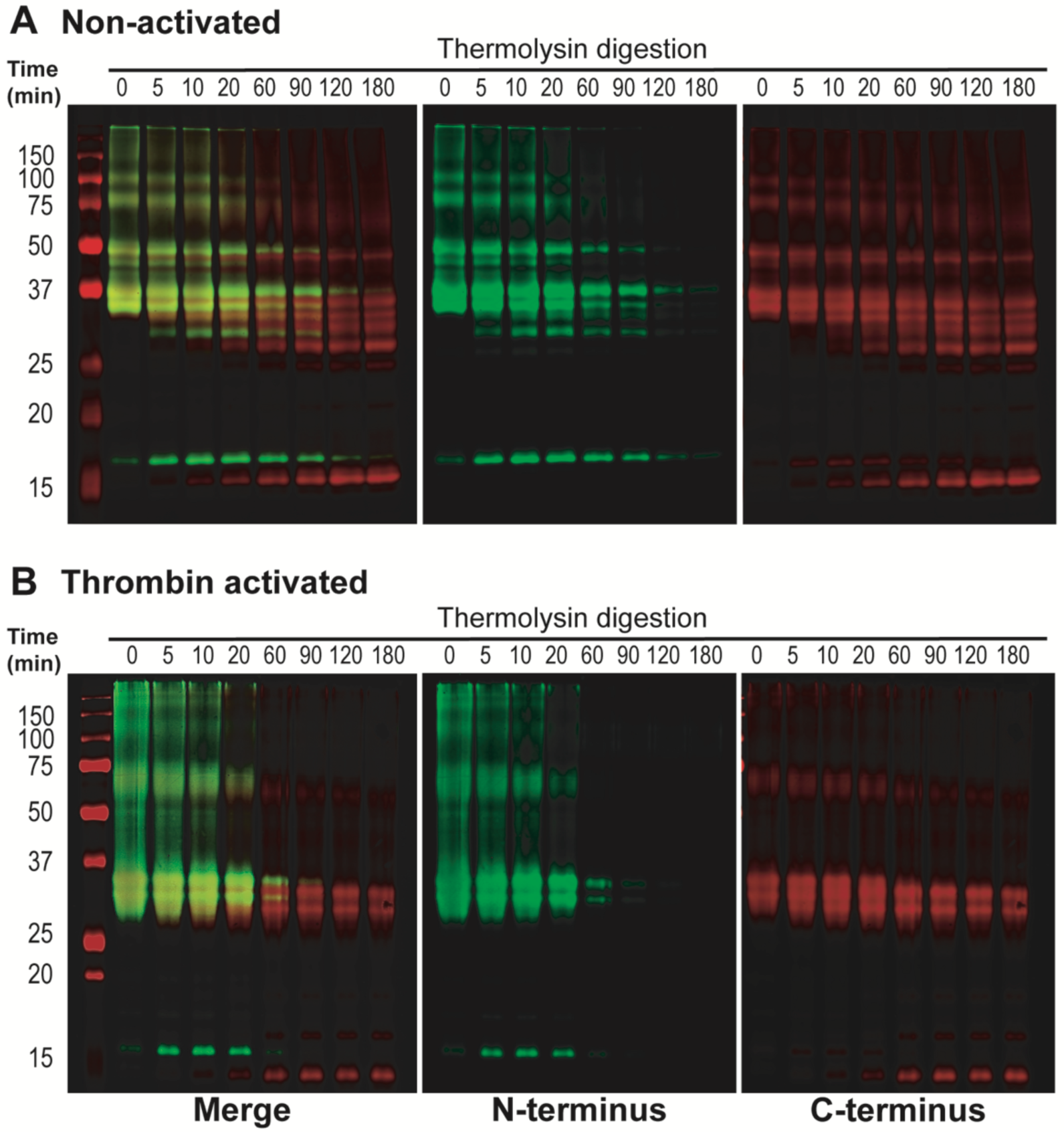
Monitoring the stability of the C- and N-terminus through Thermolysin digestion. Western Blots using two antibodies, one targeting the N-terminus (made in mouse) and the other targeting the C-terminus (made in goat) with secondary antibodies infrared fluorescent labeled with different wavelengths mouse-800 nm **(N-terminus-green)** and goat-700 nm **(C-terminus-red)**. Non-activated (A). Thrombin activated (B).

### PAR4 activation by AYPGKF

PARs can be activated independent of cleavage using soluble peptides that mimic the tethered ligand sequence.^32,33^ However, the specific response from the agonist peptides differs from that of the endogenous tethered ligand.^32,34^ We next determined the sensitivity of PAR4 to thermolysis digestion following stimulation with 1 mM of the PAR4 agonist peptide (AYPGKF), which is ∼5-fold higher than the EC_50_. The *t*_½_ of PAR4 stimulated with AYPGKF was not different from that of apo-PAR4 (Figure 5A) indicating that the agonist peptide does not induce the same conformational changes in PAR4 as thrombin. Finally, monitoring the N- and C-termini revealed that these were also unchanged compared to apo-PAR4 (Figure 5B and Figure 4A).

**Figure 5.**
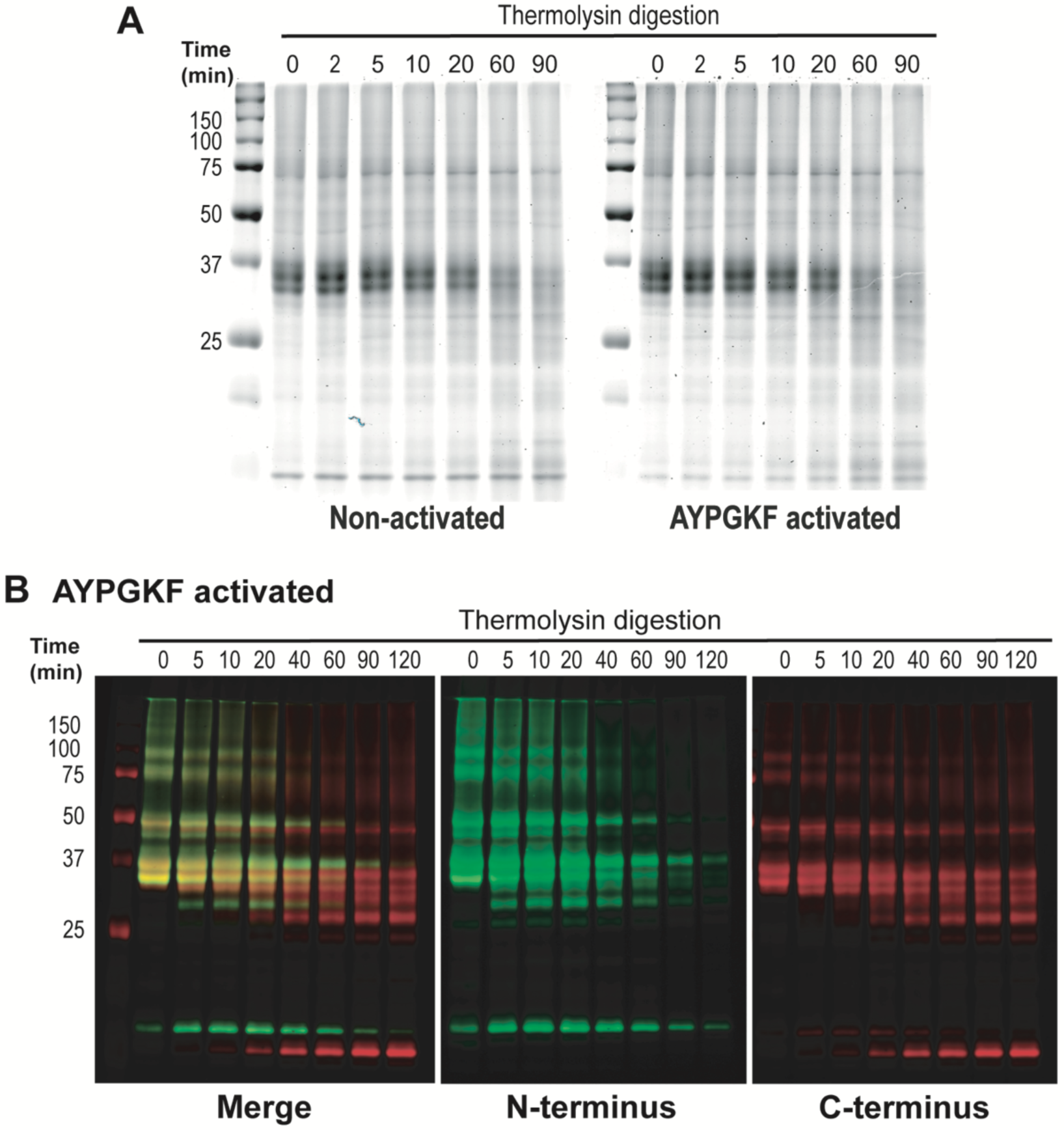
Thermolysin digestion of PAR4 non-activated and activated with AYPGKF peptide. Human PAR4 was partially digested by thermolysin as an indicator of stability; non-activated sample was used as control (A). Western Blot following the stability of the C and N terminus previously activated with AYPGKF peptide (B).

## DISCUSSION

In this report we provide two advances toward the understanding of protease activated receptor (PAR) activation mechanism. First, we describe an expression and purification strategy for full-length PAR4 without mutations or insertions to stabilize the protein. This is the first PAR to be expressed and purified with its native primary sequence that is fully activatable by an endogenous protease agonist. The second major advance builds on our purification scheme by using histidine hydrogen deuterium (His-HDX) coupled with mass spectrometry to examine the conformational dynamics of PAR4 following activation by thrombin. We compared the H/D exchange rates for each of the 9 histidines that are distributed throughout PAR4 and determined that the protein undergoes a global rearrangement following cleavage of the N-terminus by thrombin. The His-HDX data were validated by comparing the rates of proteolytic degradation by thermolysin. Cleavage of the N-terminus led to increased stability and resistance to proteolysis compared to inactive (apo) PAR4. In contrast, stimulation with the PAR4 activation peptide AYPGKF did not increase the stability of PAR4 over the apo state. Taken together, our experiments show that PAR4 undergoes a global rearrangement and becomes more stable following activation by thrombin. Finally, these studies are the first to examine PAR4 structurally and, more generally, to examine the structural dynamics of PAR activation by the tethered ligand in response to proteolytic activation.

PARs are important mediators that link extracellular proteases to intracellular signaling.^1,35^ PAR4 is the fourth member in the family that was identified and cloned in 1998.^4^ PAR4 has been primarily studied in the context of platelet signaling, which has led to efforts targeting PAR4 therapeutically.^14,15,36^ Further, polymorphisms that lead to sequence variants impact platelet reactivity in response to PAR4 agonists.^37,38^ Beyond platelets, PAR4 is also expressed on many cell types and human tissues where it has critical roles in vascular biology, inflammation, and cancer.^39–41^ Despite these numerous physiological studies, the molecular basis of PAR4 activation upon enzymatic cleavage is largely unknown. Unlike most of other receptors whose ligands are free and diffusible, the endogenous ligand of PAR4 is tethered to the protein. Therefore, PAR4 activation is likely synchronized with a global structural rearrangement of the receptor, which recruits the corresponding G-proteins that elicit downstream signaling, however this has not been directly examined experimentally.

The central question we set out to address is the structural dynamics of PAR activation in response to cleavage of the N-terminus using PAR4 activation by thrombin as a representative. Structural studies with GPCRs have largely focused on X-ray crystallography.^42^ The high resolution structures are valuable in providing detailed molecular contacts within the receptor and with specific ligands.^16,17,42^ X-ray crystallographic studies with GPCRs bound to a variety of ligands have provided a landscape of potential conformations for the receptor and how receptor allostery impacts the interaction with downstream signaling molecules.^43,44^ The primary disadvantage of structures obtained by crystallography is that no dynamic information is obtained, which is critical for PAR activation by the tethered ligand.^1^ The flexible nature of GPCRs presents a major technical challenge for crystallography studies, which requires modifications to stabilize the protein such as truncating the N- and C-termini, point mutations in the transmembrane (TM) domains, and addition of a stabilizing protein such as T4 lysozyme or BRIL.^18,42^ These modifications are at odds with determining the conformational changes of PARs following activation by a protease. To overcome this obstacle, we produced full length human PAR4 free of stabilizing sequences as a BRIL fusion protein (Figure 1A and B) in Sf9 insect cells using baculovirus.^26,18,27^ Our construct was designed to remove the FLAG epitope and BRIL sequence with TEV resulting in a final protein that begins with the first amino acid of mature human PAR4. Importantly, we show that the recombinant PAR4 is functional in response to the endogenous agonist thrombin (Figure 1). Our approach can be adapted to express other PAR family members. We anticipate that this will allow biophysical and structural experiments to determine the unique aspects of the tethered ligand for PAR1, PAR2, and PAR3.

Based on mutagenesis and domain swapping studies with PAR1, it is predicted that ECL2 is critical for PAR activation.^29^ Given that PAR1 and PAR4 have only 34% sequence identity overall and even lower in ECL2, each receptor needs to be empirically tested. The stable modifications of His residues during H/D exchange allow site specific monitoring as a protein changes conformations (Supplemental Figure 1).^19–21^ PAR4 has two histidine residues located in ECL2 (His_229_ and His_240_). Both of which had an increased *t*_½_ following activation by thrombin (Figure 2) suggesting ECL2 also has a role in PAR4 activation. Higher resolution methods are needed to determine if these are specific interactions with the tethered ligand, or if the longer *t*_½_ is due to general steric hinderance in this region. The *t*_½_ of His_380_ in the C-terminal tail increased 1.8-fold, Table 1 and Figure 2, suggesting that the C-terminus changes conformation and becomes protected from the solvent. These data are consistent with an increased stability and resistance of the C-terminus to proteolytic digestion by thermolysis (Figure 4B).

The slow H/D exchange of histidine imidazole ring, thus also its slow back exchange from D to H, allows long working times that provide the opportunity to examine proteins in complex mixtures by employing strategies that enrich for the protein of interest, such as affinity purification or immunoprecipitation, before the mass spectrometry. The ability to use these extra steps provide opportunities to examine PAR4 in its native membrane where it forms homo and heterodimers that impact it rates of activation and signaling.^24,45–48^ While advantageous for these applications, the slow His H/D exchange also creates limitations. The long exposure required for the H/D exchange on histidines greatly depends on the stability of the protein of interest. In addition, the site specific information is dependent on the number and location of His residues within the protein, however it is possible to engineer His residues at particular sites to probe regions of interest. Finally, the site specific information is useful, but the overall coverage of the protein is limited. Going forward, we can use our purification strategy to analyze PAR4 using other MS methods. By monitoring the exchange rate between the amide hydrogen at the backbone of the protein and the solvent deuterium, amide-HDX will allow us to monitor a more detailed PAR4 conformational change with a full sequence coverage and under a more physiological relevant time scale.

PARs can also be activated experimentally using soluble peptides derived from the N-terminus. For example, GYPGQV will activate PAR4 by mimicking the tethered ligand at the N-terminus following cleavage.^32,33^ However, GYPGQV is a poor PAR4 agonist, which prompted peptide library screenings for optimizing potency that AYPGKF as an alternative.^32^ The activation peptides have been essential for elucidating PAR4 specific signaling in platelets and other cells that also express PAR1, which is also activated by thrombin.^49^ However, the cellular response from the agonist peptides are not identical to thrombin suggesting that activation by the tethered ligand induces a structurally distinct conformation.^22^ We analyzed PAR4 stability by its resistance to thermolysin proteolysis following stimulation with the PAR4 agonist peptide AYPGKF. In contrast to thrombin, stimulation of PAR4 with AYPGKF (1 mM) does not increase the stability of PAR4, compare Figure 3 to Figure 5. Our His-HDX data support the hypothesis that activation by thrombin leads to a dramatic structural rearrangement of PAR4 resulting in resistance to thermolysis digest. The difference in PAR4 response is likely due to the distinct mechanisms by which the tethered ligand and the soluble activation peptide interact with the receptor.

In summary, the data presented here provide a strategy for expressing PAR4 and monitoring its activation by the tethered ligand using His-HDX. Going forward, it is now possible to further refine the purification scheme to provide a native environment using styrene maleic acid co-polymer lipid particles (SMALPS) or membrane nanodisc for applying the rapidly developing cryo-EM technologies to determine the nature of the thrombin-PAR4, PAR4-G-protein complex, or both.

## Author contributions

MM and MN conceived the study. MF, MM, and MN designed the experiments. MF, XH, MM, and MN performed the experiments and analyzed the data. MF, XH, and MN wrote the manuscript. MM critically read and edited the manuscript.

## References

1. Nieman MT. Protease-activated receptors in hemostasis. Blood. 2016;128(2):169–177.

2. Vu T-KH, Hung DT, Wheaton VI, Coughlin SR. Molecular cloning of a functional thrombin receptor reveals a novel proteolytic mechanism of receptor activation. Cell. 1991;64(6):1057–1068.

3. Zhao P, Metcalf M, Bunnett NW. Biased Signaling of Protease-Activated Receptors. Front. Endocrinol. 2014;5:.

4. Xu W-f., Andersen H, Whitmore TE, et al. Cloning and characterization of human protease-activated receptor 4. Proc. Natl. Acad. Sci. 1998;95(12):6642–6646.

5. Jacques SL, LeMasurier M, Sheridan PJ, Seeley SK, Kuliopulos A. Substrate-Assisted Catalysis of the PAR1 Thrombin Receptor: ENHANCEMENT OF MACROMOLECULAR ASSOCIATION AND CLEAVAGE. J. Biol. Chem. 2000;275(52):40671–40678.

6. Myles T, Le Bonniec BF, Stone SR. The dual role of thrombin’s anion-binding exosite-I in the recognition and cleavage of the protease-activated receptor 1: Kinetics for the cleavage of PAR1 by thrombin. Eur. J. Biochem. 2001;268(1):70–77.

7. Jacques SL, Kuliopulos A. Protease-activated receptor-4 uses dual prolines and an anionic retention motif for thrombin recognition and cleavage. Biochem. J. 2003;376(3):733–740.

8. Nieman MT, Schmaier AH. Interaction of Thrombin with PAR1 and PAR4 at the Thrombin Cleavage Site †. Biochemistry. 2007;46(29):8603–8610.

9. Nieman MT. Protease-Activated Receptor 4 Uses Anionic Residues To Interact with α-Thrombin in the Absence or Presence of Protease-Activated Receptor 1 †. Biochemistry. 2008;47(50):13279–13286.

10. Duvernay MT, Temple KJ, Maeng JG, et al. Contributions of Protease-Activated Receptors PAR1 and PAR4 to Thrombin-Induced GPIIbIIIa Activation in Human Platelets. Mol. Pharmacol. 2016;91(1):39–47.

11. Covic L, Gresser AL, Kuliopulos A. Biphasic kinetics of activation and signaling for PAR1 and PAR4 thrombin receptors in platelets. Biochemistry. 2000;39(18):5458–5467.

12. Shapiro MJ, Weiss EJ, Faruqi TR, Coughlin SR. Protease-activated Receptors 1 and 4 Are Shut Off with Distinct Kinetics after Activation by Thrombin. J. Biol. Chem. 2000;275(33):25216–25221.

13. Han X, Nieman MT. PAR4 (Protease-Activated Receptor 4): PARticularly Important 4 Antiplatelet Therapy. Arterioscler. Thromb. Vasc. Biol. 2018;38(2):287–289.

14. Wong PC, Seiffert D, Bird JE, et al. Blockade of protease-activated receptor-4 (PAR4) provides robust antithrombotic activity with low bleeding. Sci. Transl. Med. 2017;9(371):eaaf5294.

15. Wilson SJ, Ismat FA, Wang Z, et al. PAR4 (Protease-Activated Receptor 4) Antagonism With BMS-986120 Inhibits Human Ex Vivo Thrombus Formation. Arterioscler. Thromb. Vasc. Biol. 2018;38(2):448–456.

16. Zhang C, Srinivasan Y, Arlow DH, et al. High-resolution crystal structure of human protease-activated receptor 1. Nature. 2012;492(7429):387–392.

17. Cheng RKY, Fiez-Vandal C, Schlenker O, et al. Structural insight into allosteric modulation of protease-activated receptor 2. Nature. 2017;545(7652):112–115.

18. Chun E, Thompson AA, Liu W, et al. Fusion Partner Toolchest for the Stabilization and Crystallization of G Protein-Coupled Receptors. Structure. 2012;20(6):967–976.

19. Lodowski DT, Palczewski K, Miyagi M. Conformational Changes in the G Protein-Coupled Receptor Rhodopsin Revealed by Histidine Hydrogen−Deuterium Exchange. Biochemistry. 2010;49(44):9425–9427.

20. Miyagi M, Wan Q, Ahmad MdF, et al. Histidine Hydrogen-Deuterium Exchange Mass Spectrometry for Probing the Microenvironment of Histidine Residues in Dihydrofolate Reductase. PLoS ONE. 2011;6(2):e17055.

21. Mullangi V, Zhou X, Ball DW, Anderson DJ, Miyagi M. Quantitative Measurement of the Solvent Accessibility of Histidine Imidazole Groups in Proteins. Biochemistry. 2012;51(36):7202–7208.

22. McLaughlin JN, Shen L, Holinstat M, et al. Functional Selectivity of G Protein Signaling by Agonist Peptides and Thrombin for the Protease-activated Receptor-1. J. Biol. Chem. 2005;280(26):25048–25059.

23. Mumaw MM, de la Fuente M, Arachiche A, Wahl JK, Nieman MT. Development and characterization of monoclonal antibodies against Protease Activated Receptor 4 (PAR4). Thromb. Res. 2015;135(6):1165–1171.

24. Smith TH, Coronel LJ, Li JG, et al. Protease-activated Receptor-4 Signaling and Trafficking Is Regulated by the Clathrin Adaptor Protein Complex-2 Independent of β-Arrestins. J. Biol. Chem. 2016;291(35):18453–18464.

25. Gulati S, Jastrzebska B, Banerjee S, et al. Photocyclic behavior of rhodopsin induced by an atypical isomerization mechanism. Proc. Natl. Acad. Sci. 2017;114(13):E2608–E2615.

26. Chu R, Takei J, Knowlton JR, et al. Redesign of a Four-helix Bundle Protein by Phage Display Coupled with Proteolysis and Structural Characterization by NMR and X-ray Crystallography. J. Mol. Biol. 2002;323(2):253–262.

27. Liu W, Chun E, Thompson AA, et al. Structural Basis for Allosteric Regulation of GPCRs by Sodium Ions. Science. 2012;337(6091):232–236.

28. Jahnsen JA, Uhlén S. The N-terminal region of the human 5-HT2C receptor has as a cleavable signal peptide. Eur. J. Pharmacol. 2012;684(1-3):44–50.

29. Nanevicz T, Ishii M, Wang L, et al. Mechanisms of Thrombin Receptor Agonist Specificity: CHIMERIC RECEPTORS AND COMPLEMENTARY MUTATIONS IDENTIFY AN AGONIST RECOGNITION SITE. J. Biol. Chem. 1995;270(37):21619–21625.

30. Hubbard SJ. The structural aspects of limited proteolysis of native proteins. Biochim. Biophys. Acta BBA - Protein Struct. Mol. Enzymol. 1998;1382(2):191–206.

31. Fontana A, de Laureto PP, Spolaore B, et al. Probing protein structure by limited proteolysis. 2004;51:23.

32. Faruqi TR, Weiss EJ, Shapiro MJ, Huang W, Coughlin SR. Structure-Function Analysis of Protease-activated Receptor 4 Tethered Ligand Peptides: DETERMINANTS OF SPECIFICITY AND UTILITY IN ASSAYS OF RECEPTOR FUNCTION. J. Biol. Chem. 2000;275(26):19728–19734.

33. Hollenberg MD, Saifeddine M. Proteinase-activated receptor 4 (PAR4): activation and inhibition of rat platelet aggregation by PAR4-derived peptides. 2001;79:6.

34. Hollenberg MD. Proteinase-activated receptors: Tethered ligands and receptor-activating peptides. Drug Dev. Res. 2003;59(4):336–343.

35. Sidhu T, French S, Hamilton J. Differential Signaling by Protease-Activated Receptors: Implications for Therapeutic Targeting. Int. J. Mol. Sci. 2014;15(4):6169–6183.

36. Han X, Nieman MT. PAR4 (Protease-Activated Receptor 4): PARticularly Important 4 Antiplatelet Therapy. Arterioscler. Thromb. Vasc. Biol. 2018;38(2):287–289.

37. Edelstein LC, Simon LM, Lindsay CR, et al. Common variants in the human platelet PAR4 thrombin receptor alter platelet function and differ by race. Blood. 2014;124(23):3450–3458.

38. Whitley MJ, Henke DM, Ghazi A, et al. The protease-activated receptor 4 Ala120Thr variant alters platelet responsiveness to low-dose thrombin and protease-activated receptor 4 desensitization, and is blocked by non-competitive P2Y 12 inhibition. J. Thromb. Haemost. 2018;16(12):2501–2514.

39. Reed CE, Kita H. The role of protease activation of inflammation in allergic respiratory diseases. J. Allergy Clin. Immunol. 2004;114(5):997–1008.

40. Lee H, Hamilton JR. Physiology, pharmacology, and therapeutic potential of protease-activated receptors in vascular disease. Pharmacol. Ther. 2012;134(2):246–259.

41. Wojtukiewicz MZ, Hempel D, Sierko E, Tucker SC, Honn KV. Thrombin—unique coagulation system protein with multifaceted impacts on cancer and metastasis. Cancer Metastasis Rev. 2016;35(2):213–233.

42. Cherezov V, Abola E, Stevens RC. Recent Progress in the Structure Determination of GPCRs, a Membrane Protein Family with High Potential as Pharmaceutical Targets. Membr. Protein Struct. Determ. 2010;654:141–168.

43. Katritch V, Cherezov V, Stevens RC. Structure-Function of the G Protein–Coupled Receptor Superfamily. Annu. Rev. Pharmacol. Toxicol. 2013;53(1):531–556.

44. Luttrell LM, Maudsley S, Bohn LM. Fulfilling the Promise of “Biased” G Protein–Coupled Receptor Agonism. Mol. Pharmacol. 2015;88(3):579–588.

45. Arachiche A, Mumaw MM, de la Fuente M, Nieman MT. Protease-activated Receptor 1 (PAR1) and PAR4 Heterodimers Are Required for PAR1-enhanced Cleavage of PAR4 by α-Thrombin. J. Biol. Chem. 2013;288(45):32553–32562.

46. Arachiche A, de la Fuente M, Nieman MT. Calcium Mobilization And Protein Kinase C Activation Downstream Of Protease Activated Receptor 4 (PAR4) Is Negatively Regulated By PAR3 In Mouse Platelets. PLoS ONE. 2013;8(2):e55740.

47. Khan A, Li D, Ibrahim S, Smyth E, Woulfe DS. The Physical Association of the P2Y12 Receptor with PAR4 Regulates Arrestin-Mediated Akt Activation. Mol. Pharmacol. 2014;86(1):1–11.

48. Li D, D’Angelo L, Chavez M, Woulfe DS. Arrestin-2 Differentially Regulates PAR4 and ADP Receptor Signaling in Platelets. J. Biol. Chem. 2011;286(5):3805–3814.

49. Kahn ML, Nakanishi-Matsui M, Shapiro MJ, Ishihara H, Coughlin SR. Protease-activated receptors 1 and 4 mediate activation of human platelets by thrombin. J. Clin. Invest. 1999;103(6):879–887.

