## Supplemental Data for "Expression and purification of Protease Activated Receptor 4 (PAR4) and analysis with histidine hydrogen deuterium exchange"

**Supplemental Figure 1. Schematic view of the Hydrogen-Deuterium exchange**. A schematic showing the chemical exchange of the hydrogen at the C2 carbon on the imidazole ring of histidine residues (A). The back exchange is slow at this position allowing for long time scale measurements using histidine residues as site specific probes within a protein. Histidine hydrogen deuterium exchange (His-HDX) mass spectrometry takes advantage of this property of the C2 carbon by incubating the protein of interest in $D_2O$ for 72 hours. During this incubation, hydrogens are exchanged to deuterium. The amide hydrogens quickly back exchange leaving deuterium atoms that are stably incorporated into the C2 carbon of histidines within the protein (B).

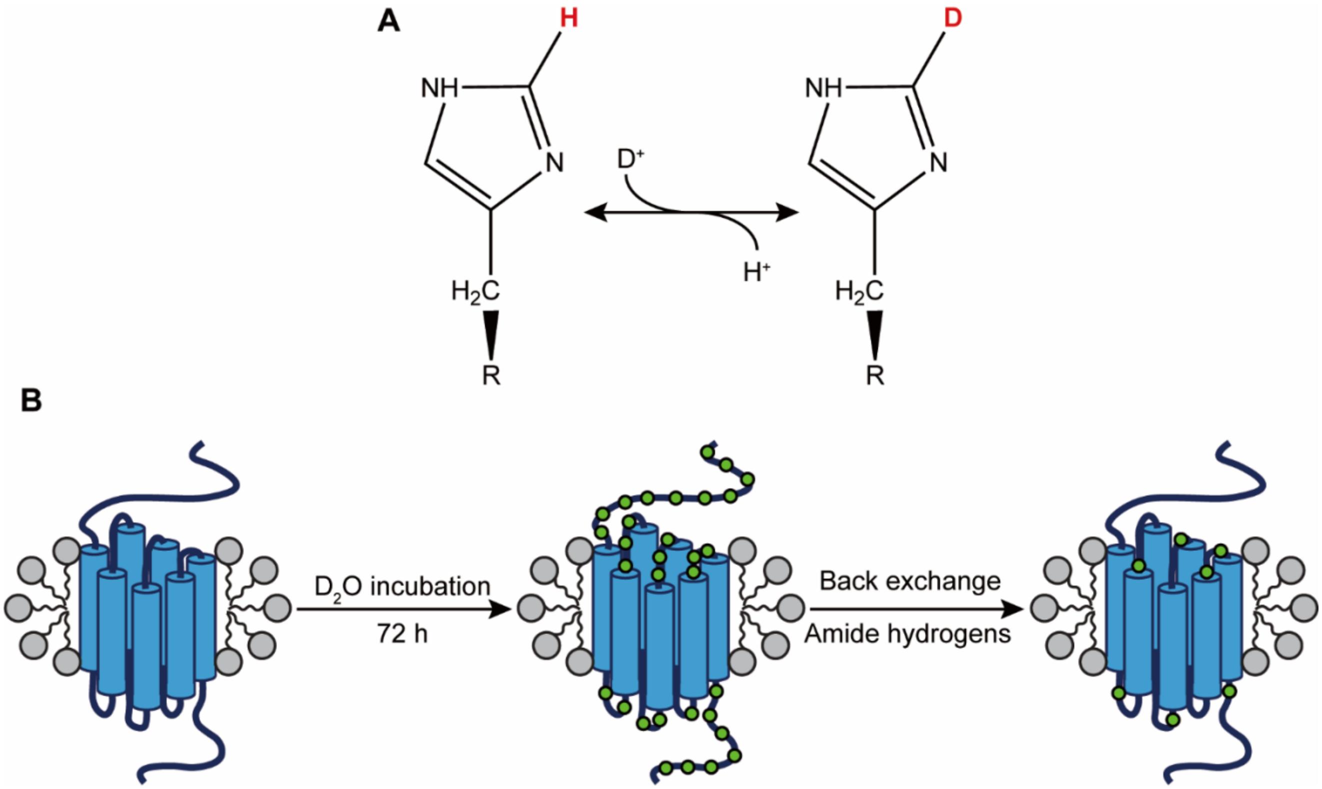

**Supplemental Figure 2. Histidine-peptide coverage of His-HDX experiment.** A snake diagram of PAR4 showing the coverage and the location of the peptides that contain histidines analyzed by mass spectrometry. Each peptide contained a single His residue. Note, the peptides in ECL2 are adjacent to one another ($C_{228}$HDALPLD and $A_{236}$QASHW).

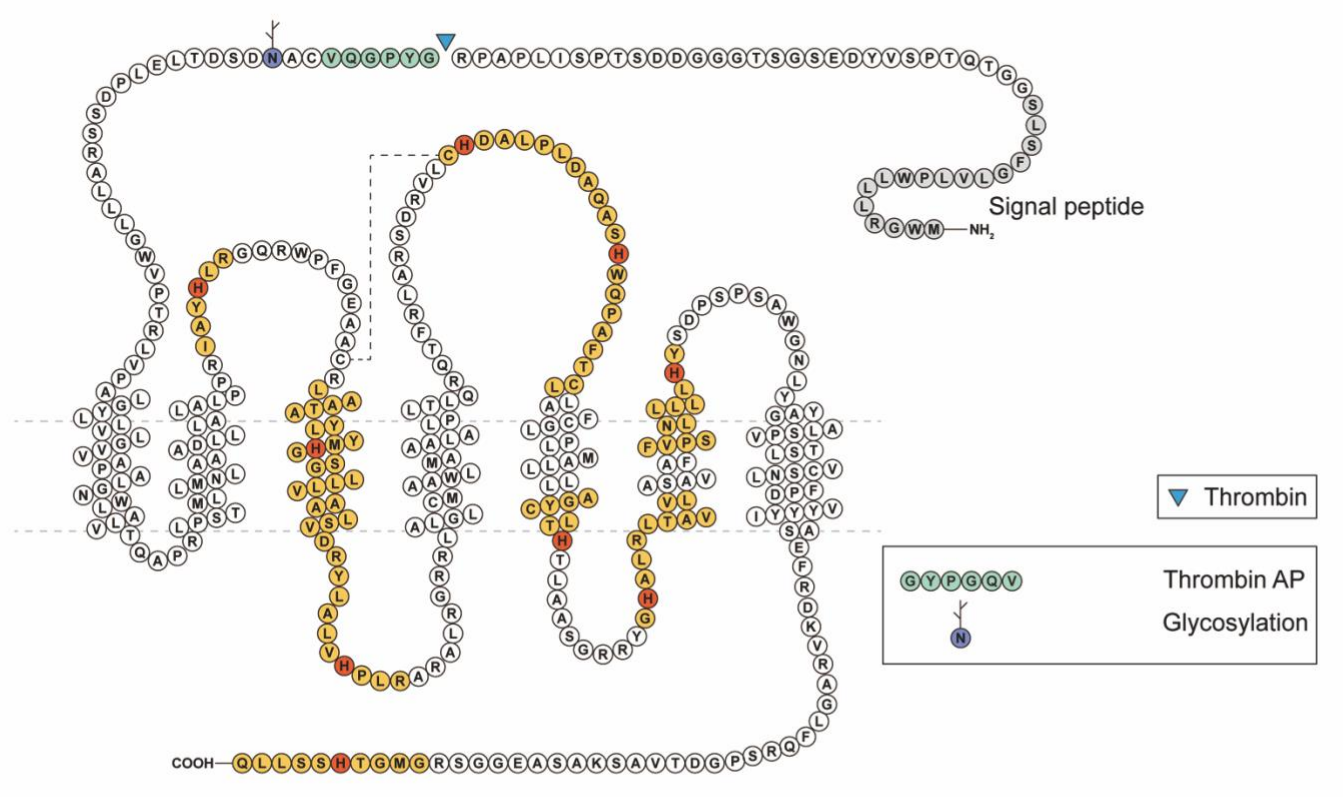

**Supplemental Figure 3. Mapping the epitope of PAR4 C-20 antibody.** Lysates of HEK293 cells expressing WT-PAR4 and two C–terminal truncation mutants were blotted with the anti-PAR4 C-20 antibody. α-Actinin is used as a loading control. The decreased intensity in the C-20 antibody interacts with the last 12 amino acids of PAR4 at the very end of the C-terminus, more specifically between $Gly_{373}$ and the C-terminus (A). A diagram indicating the site of the truncation PAR4 (B). The detailed sequence of PAR4 is shown in Supplemental Figure 2.

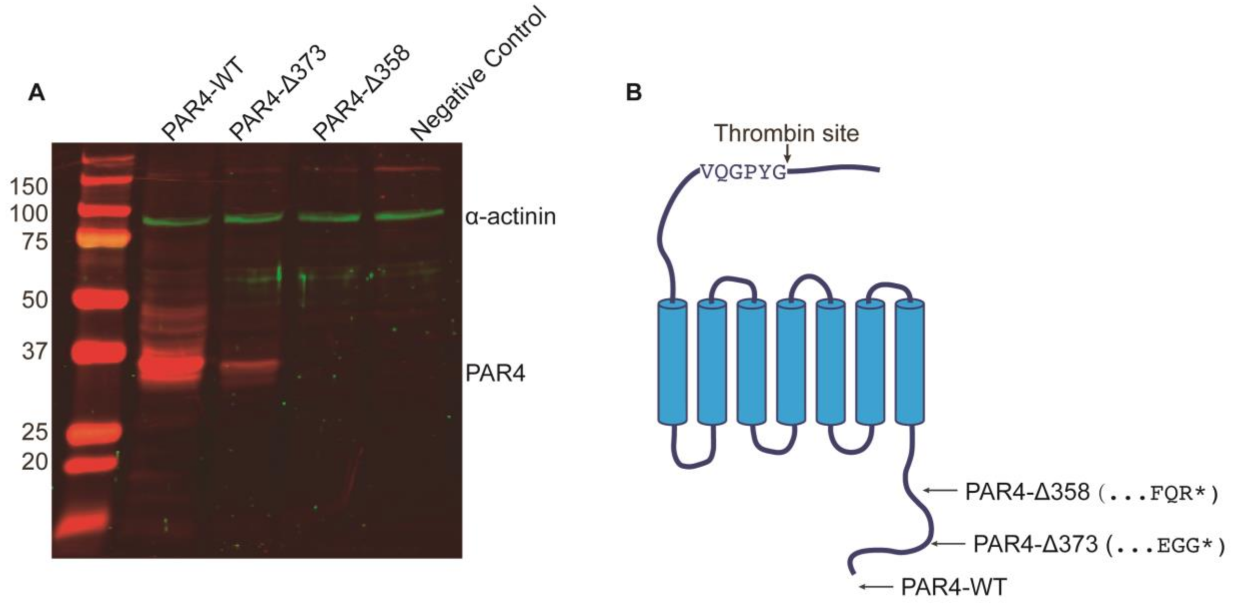

**Supplemental Table1**.

| Supplemental Table 1. Histidine peptides analyzed from PAR4 digested with trypsin or chymotrypsin | |
|---|---|
| $His_{136}$ | RIAY**H**LRG |
| $His_{159}$ | RLATAALYG**H**MYGSVLLLAAVSLDRY |
| $His_{159}$ | YG**H**MYGSVLL |
| His $_{180}$ | RYLALV**H**PLRA |
| $His_{380}$ | RGMGT**H**SSLLQ |
| $His_{159}$ | YG**H**MYGSVLL |
| $His_{180}$ | LALV**H**PLR |
| $His_{229}$ | LC**H**DALPLD |
| $His_{240}$ | AQAS**H**W |
| $His_{269}$ | LCYGATL**H**TLA |
| $His_{280}$ | G**H**ALRLTAVVL |
| $His_{306}$ | LL**H**YSDPSPSAWG |
| $His_{380}$ | MGT**H**SSLLQ |
